# A comparative recombination analysis of human coronaviruses and implications for the SARS-CoV-2 pandemic

**DOI:** 10.1101/2021.03.07.434287

**Authors:** Simon Pollett, Matthew A Conte, Mark Sanborn, Richard G Jarman, Grace M. Lidl, Kayvon Modjarrad, Irina Maljkovic Berry

## Abstract

The SARS-CoV-2 pandemic prompts evaluation of recombination in human coronavirus (hCoV) evolution. We undertook recombination analyses of 158,118 public seasonal hCoV, SARS-CoV-1, SARS-CoV-2 and MERS-CoV genome sequences using the RDP4 software. We found moderate evidence for 8 SARS-CoV-2 recombination events, two of which involved the spike gene, and low evidence for one SARS-CoV-1 recombination event. Within MERS-CoV, 229E, OC43, NL63 and HKU1 datasets, we noted 7, 1, 9, 14, and 1 high-confidence recombination events, respectively. There was propensity for recombination breakpoints in structural genes, and recombination severely skewed the temporal structure of these data, especially for NL63 and OC43. Bayesian time-scaled analyses on recombinant-free data indicated the sampled diversity of seasonal CoVs emerged in the last 70 years, with 229E displaying continuous lineage replacements. These findings emphasize the importance of genomic based surveillance to detect recombination in SARS-CoV-2, particularly if recombination may lead to immune evasion.

## INTRODUCTION

The emergence of SARS-CoV-2 has generated interest in role of recombination in the evolution of this and other human coronaviruses (hCoV). Recombination has been observed in many RNA viruses and is noted to occur at a higher frequency in positive-sense RNA viruses, a category that includes SARS-CoV-2 and other medically important coronaviruses (1–3). From an evolutionary biology perspective, it remains unclear why recombination occurs in RNA viruses. Hypotheses include recombination being an incidental outcome of RNA polymerase function, or an evolutionary favorable purge of deleterious genotypes and/or generation of advantageous genotypes (1).

While several studies have examined the putative role of recombination in the zoonotic emergence of SARS-CoV-2 (4–6), few have focused on the emergence of recombination during the first year of the SARS-CoV-2 pandemic. To date, none have systematically examined the genetic propensity of recombination across all human coronaviruses in order to predict the possible evolutionary future of SARS-CoV-2, despite prior observations of recombination in OC43-hCoV, HKU1-hCoV, NL63-hCoV, and MERS-CoV (7–19).

RNA virus recombination has been associated with changes in host range, host response and virulence (1). Identifying the presence of recombination, or predicting the risk of recombination, in viral populations of SARS-CoV-2 is critical for several reasons. First, circulating recombinants may complicate molecular diagnostic performance. Second, recombinants may cause rapid escape from naturally acquired immunity, as has been observed in the norovirus genus, which has caused pandemics due to the rapid emergence of new genotypes generated by recombination of structural genes (20). For SARS-CoV-2, such an event may have major implications, especially if circulating recombinant results in escape from both natural and vaccine induced immunity (21). Finally, genomic epidemiology has increasingly been shown to be an important public health tool for SARS-CoV-2 and failing to accommodate for recombinant data may lead to incorrect epidemiological inference because of possible phylogenetic incongruence (22, 23).

We undertook a comparative recombination analysis of all published SARS-CoV-1, SARS-CoV-2, MERS-CoV and seasonal hCoV (OC43, 229E, NL63 and HKU1) genomes. Specifically, we aimed to identify the frequency and genomic location of recombination events in these hCoV types, and estimate the impact of these recombinants on hCoV emergence dates (TMRCA). As part of this analysis, we reconstructed the time-scale of circulating seasonal hCoV evolution and lineage replacement to provide insights into the possible future evolutionary trajectory of SARS-CoV-2.

## RESULTS

### There is moderate evidence for recombination in several SARS-CoV-2 genomes through October 2020

Among SARS-CoV-2 genomes, we detected a total of 8 recombination events detected by at least three detection methods, though these events were possibly caused by another non-recombinant process (Table 1). Some of these events, while not supported by a high level of evidence, were noted in multiple sequences and may therefore represent circulating recombinant forms (Supplemental Table 1). However, these recombination events were not found across all subsampled datasets. For those recombination events with moderate evidence, we noted that half of the events (4/8) comprised breakpoints in the structural genes, and one quarter (2/8) occurred within the spike gene (Supplemental Table 2).

**Table 1.** Frequency of recombination events detected in 229E, NL63, OC43, HKU1, MERS-CoV, SARS-CoV1 and SARS-CoV-2, stratified by level of evidence

| Coronavirus species | *n* genome sequences | Recombination events detected by any method | Recombination events detected by $\geq 3$ methods | Recombination events detected by $\geq 3$ methods and without another evolutionary process possibly explaining the recombination signal (high evidence) | Recombination events with high level evidence seen in multiple genomes |
| --- | --- | --- | --- | --- | --- |
| hCoV-229E | 22 | 4 | 3 | 1 | 1 |
| hCoV-NL63 | 65 | 31 | 24 | 14 | 7 |
| hCoV-OC43 | 138 | 23 | 16 | 9 | 6 |
| hCoV-HKU1 | 37 | 14 | 9 | 1 | 1 |
| MERS-CoV | 365 | 12 | 10 | 7 | 6 |
| SARS-CoV-1 | 49 | 1 | 0 | 0 | 0 |
| SARS-CoV-2 | 100296<sup>a</sup> | 33 | 8 | 0 | 0 |

<sup>a</sup>Randomly subsampled into 100 x n = 300 independent datasets

### Recombination is relatively frequent in seasonal endemic coronaviruses and has a propensity for the structural genes

Within the 229E, OC43, NL63 and HKU1 datasets, we noted 1, 9, 14, and 1 high confidence recombination events, respectively (Table 1). These recombination events were found in 13.6%, 20.3%, 89.2%, and 35.1% of the analyzed sequences, respectively. In the OC43 and NL63 datasets, we noted significantly more breakpoints in the structural genes than in the non-structural genes (p = 0.0004, p = 0.0032 respectively) (Figure 1).

**Figure 1.**
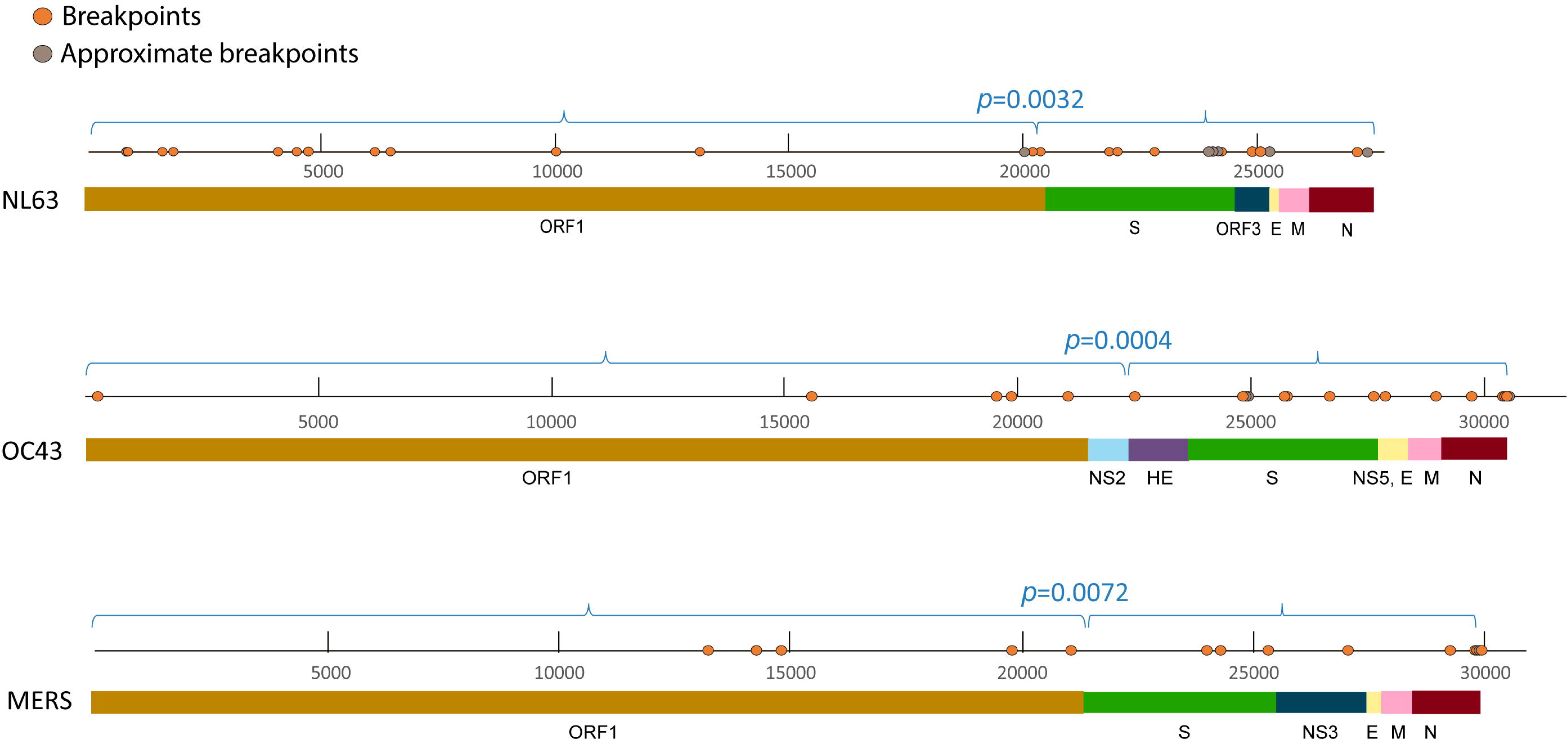
Estimated recombination breakpoint positions NL63, OC43 and MERS-CoV whole genomes. P-values for the frequency of recombination breakpoints in structural versus non-structural genes are derived by the χ^2^ test. Approximate breakpoints are breakpoints that could not be placed with certainty due to overlapping recombination or other reasons.

Recombinants were noted across entire clades of HKU1, NL63 and OC43 viruses, with clusters of genomes sharing identical recombination patterns, indicating spread of a recombinant virus following its emergence through a recombination event. The OC43 and NL63 datasets also contained genomes with unique recombination patterns (singletons) (Figure 2, Figure 3). Furthermore, we noticed some singletons falling within already recombinant clades, indicating presence of successful second generation recombination (Figure 2, Figure 3). A 10^th^ OC43 event with moderate recombination evidence based on RDP4 results showed topological incongruence in subgenomic phylogenetic trees and was therefore removed for further time-scaled analyses.

**Figure 2.**
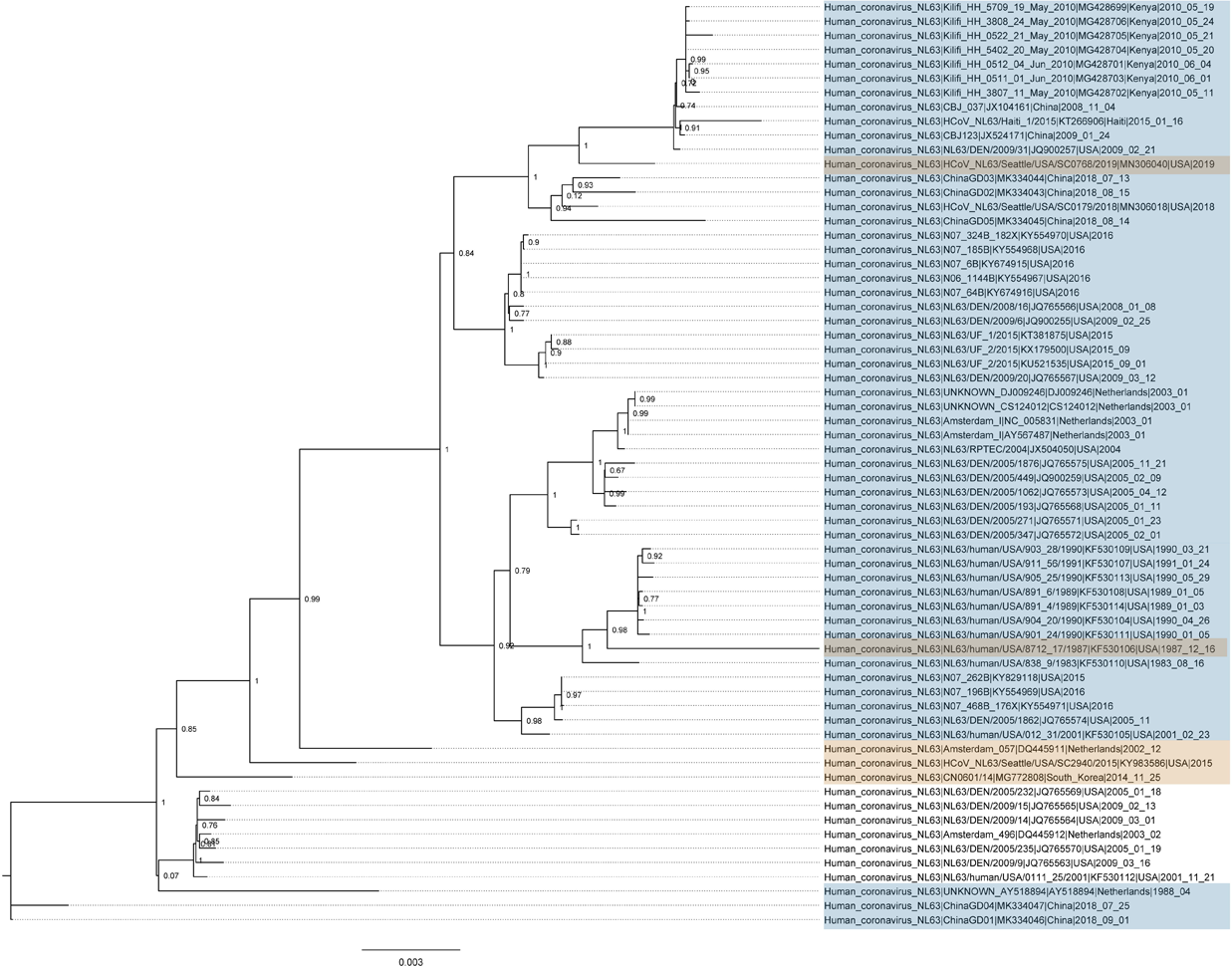
Maximum likelihood phylogeny of recombinants in NL63. Scale represents nucleotides per site. Recombinant events with multiple genomes are marked in blue, or as singletons are marked in yellow. Phylogeny was rooted with a 229E outgroup (removed for clarity).

**Figure 3.**
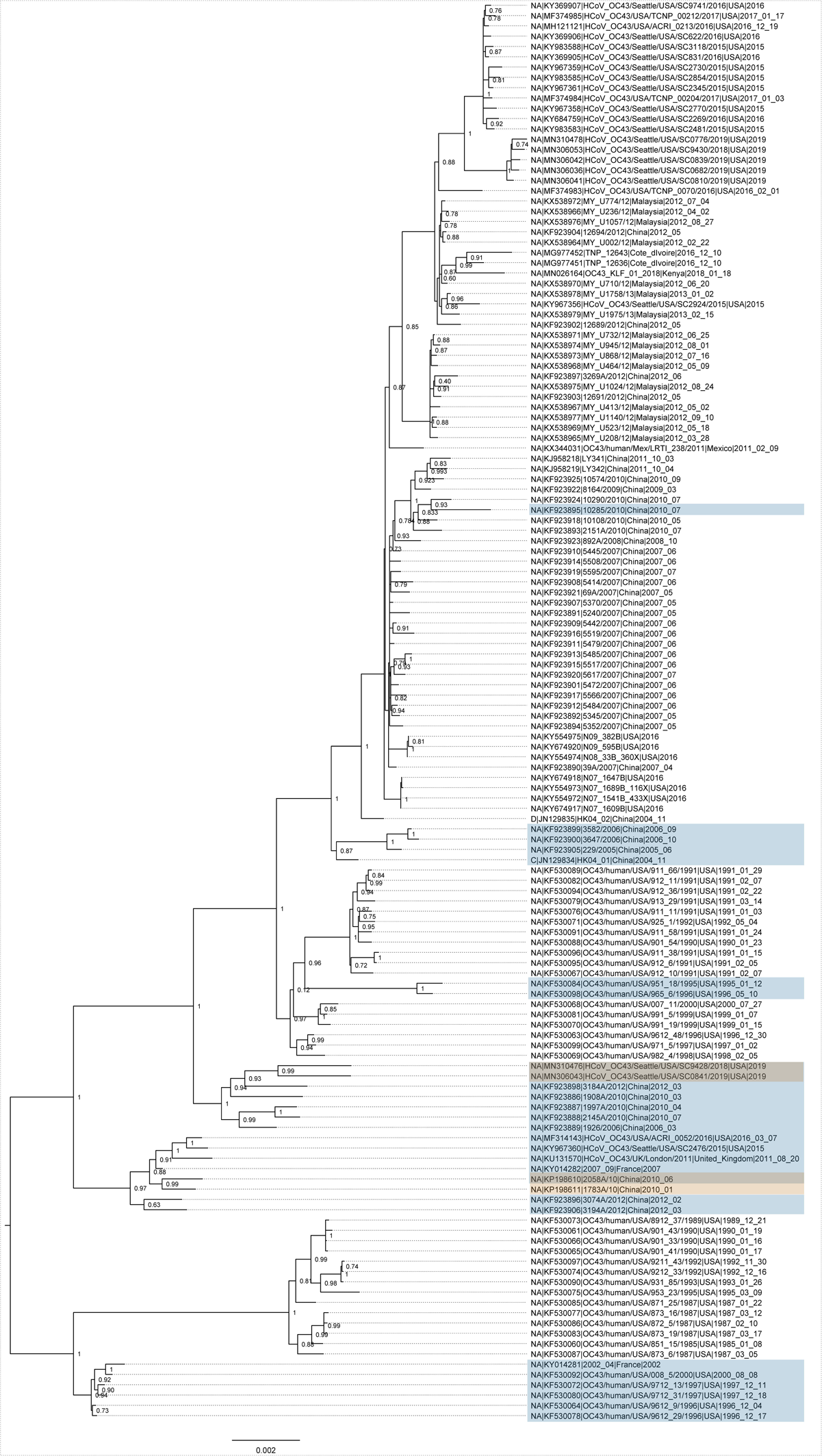
Maximum likelihood phylogenies of recombinants in OC43. Scale represents nucleotides per site. Recombinant events with multiple genomes are marked in blue, or as singletons are marked in yellow. Phylogeny was rooted with an HKU1 outgroup (removed for clarity).

### MERS-CoV but not SARS-CoV-1 is characterized by frequent recombination with a propensity for structural genes

Within the MERS-CoV dataset, we detected 7 recombination events with a high level of confidence (detected by 3 or more methods and not explained by another evolutionary process) (Table 1). These recombination events were found in 14.5% of all analyzed MERS-CoV genomes. Of these, 6 were found across clades, suggesting recombinants were sufficiently fit for onward transmission (Figure 4). We noted recombinant clades defined by camel hosts, as well as camel and human hosts (Figure 4), suggestive of inter-host recombinant spread. Moreover, we noted significantly more breakpoints in the structural genes compared to the non-structural genes (p < 0.001) (Figure 1). We noted only a single low evidence recombination event involving the structural region of the SARS-CoV1 genome (4503 −25998 nt), but this was not confirmed by multiple methods (Table 1).

**Figure 4.**
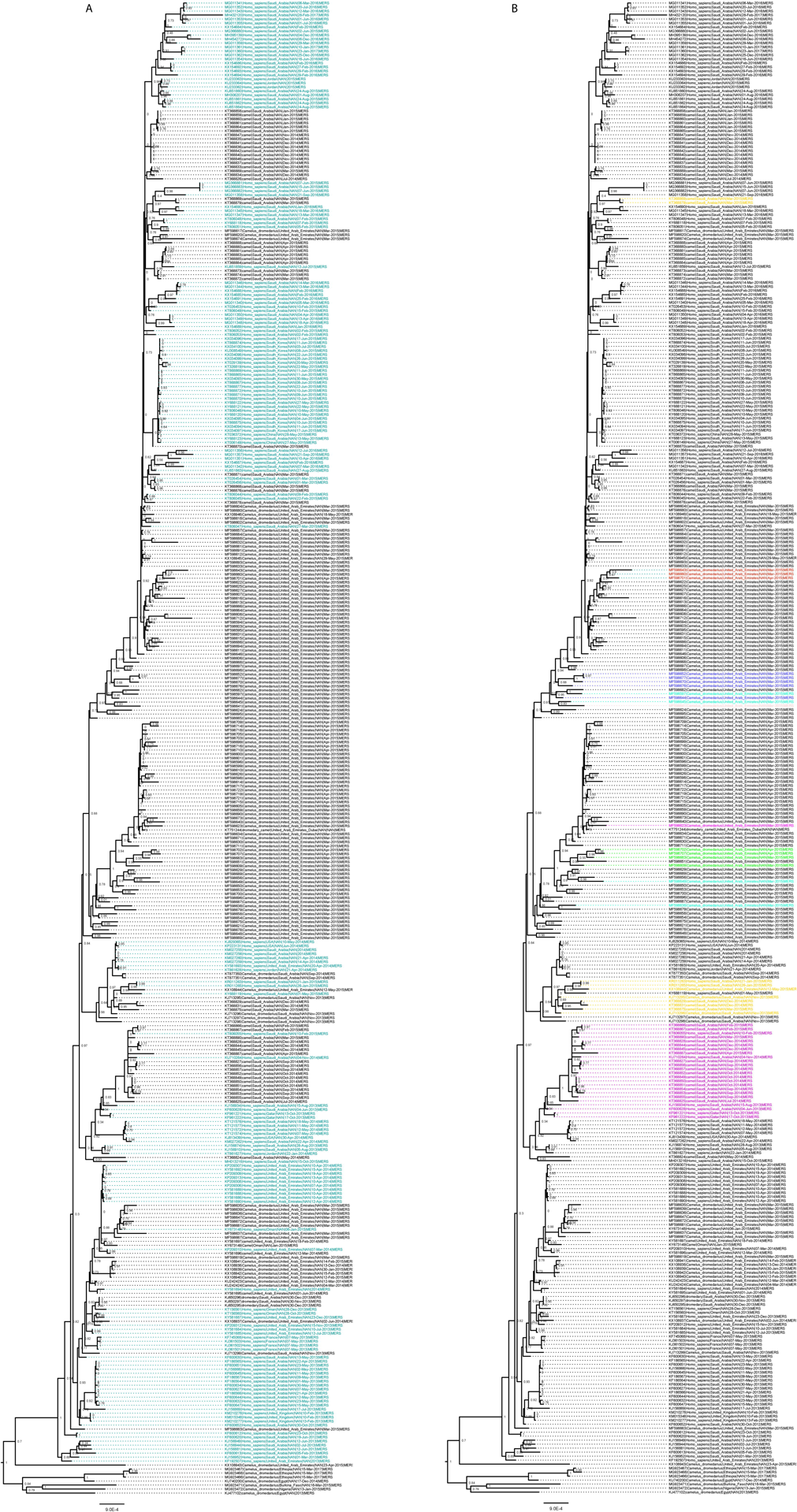
Maximum likelihood phylogeny of recombinants in MERS-CoV. Scale represents nucleotides per site. A) Taxa colored by host (camel = black, human = green). B) Colored taxa indicate confirmed recombinant clades.

### Recombination in seasonal coronaviruses substantially alters estimates of temporal structure

Our approach to identification of recombinants and their subsequent removal from the datasets led to major improvements in estimated temporal structure of hCoV-OC43, hCoV-NL63 and hCoV-HKU1. These changes involved both the regression coefficient and the regression intercept, which serve as crude estimates of evolutionary rates and TMRCA, and are often used to assess the clock-likeness (linear relationship of genetic distance across sequence sampling times) of the data for further analyses (Table 2). In contrast, removal of the single 229E recombinant did not cause substantive change in estimated temporal structure as estimated by regression coefficient and TMRCA (Table 2).

**Table 2.** Root-to-tip regression coefficient and intercept of seasonal hCoV phylogenies with and without recombinants removed

| Recombinants not removed |  |  |  |  |
| --- | --- | --- | --- | --- |
| Lineage | Sequences (n) | Date range (years) | Slope coefficient<sup>a</sup> | Intercept (TMRCA)<sup>b</sup> |
| hCoV-229E | 22 | 26.36 | $2.69 \times 10^{-4}$ | 1990 A.D |
| hCoV-NL63 | 65 | 35.38 | $-1.00 \times 10^{-4}$ | 2149 A.D |
| hCoV-OC43 | 138 | 33.98 | -0.00 | 27359 A.D |
| hCoV-HKU1 | 37 | 14.17 | $4.52 \times 10^{-4}$ | 1941 A.D |
| Recombinants removed |  |  |  |  |
| Lineage | Sequences (n) | Date range (years) | Slope coefficient<sup>a</sup> | Intercept<sup>b</sup> |
| hCoV-229E | 19 | 26.36 | $2.65 \times 10^{-4}$ | 1990 A.D |
| hCoV-NL63* | 56 | 35.04 | $7.68 \times 10^{-5}$ | 1944 A.D |
| hCoV-OC43 | 110 | 34.00 | $2.83 \times 10^{-4}$ | 1967 A.D |
| hCoV-HKU1 | 24 | 13.41 | $1.00 \times 10^{-3}$ | 1978 A.D |

<sup>a</sup>Approximates evolutionary rate (substitutions/site/year)
<sup>b</sup>Approximates TMRCA
\*Non-recombinant region was used, with genomes containing breakpoints in this region removed (N=9)

### The current global diversity of seasonal hCoVs arose across the last 70 years

We estimated a TMRCA date (year A.D) of 1989, 1970, 1964, and 1951 for the 229E, OC43, NL63 and HKU1 lineages, respectively, indicating relatively recent emergence of the current seasonal hCoV lineages (Table 3, Supplemental Figures 1-4). Importantly, these do not necessarily represent *de novo* emergence of these viruses from animal origins but rather may represent the divergence from older OC43, NL63 and HKU1 lineages, respectively. Indeed, the TMRCA for 229E is preceded by historical descriptions of the circulation of this virus in humans, which was discovered in 1965 (24). We therefore extended our 229E full genome analysis and inferred TMRCA estimates from partial genome datasets of the complete N gene (N=102, length=1167nt), the complete S gene (Supplemental Figure 5) (N=78, length=3522nt), the RBD (S1) domain S gene (N=90, length=1650nt), and concatenated S and N genes (N=64, length=4689nt). These yielded TMRCA point estimates between 1964 and 1969 (229E RBD S1 gene TMRCA = 1965.6; 229E S gene TMRCA = 1964.8; 229E N gene TMRCA = 1969.1; 229E N and S gene TMRCA = 1964.0).

**Table 3.** Bayesian TMRCA estimates for 229E, HKU1, NL63 and OC43^a^

|  | TMRCA<br>(A.D) | Lower<br>95% HPD | Upper<br>95% HPD | Nucleotide<br>Subst Model | Clock model | Demographic<br>model |
| --- | --- | --- | --- | --- | --- | --- |
| 229E<sup>a</sup> | 1989 | 1988 | 1990 | SYM+I+G | Strict | Constant |
| HKU1<sup>a</sup> | 1951 | 1842 | 1998 | GTR+G+I | UCLN | Bayesian Skyline |
| NL63<sup>b</sup> | 1964 | 1945 | 1978 | HKY+I | UCLN | Bayesian Skyline |
| OC43<sup>a</sup> | 1970 | 1960 | 1978 | GTR+I | UCLN | Bayesian Skyline |

<sup>a</sup>Recombinant genomes removed
<sup>b</sup>Non-recombinant region 13093-20198
UCLN = uncorrelated lognormal; TMRCA = time to most recent common ancestor

## DISCUSSION

We performed a comprehensive recombination analysis across all medically important human coronaviruses, with an overarching aim to identify the current and future risk of recombinant emergence in the SARS-CoV-2 pandemic. We note moderate evidence for SARS-CoV-2 recombination during the first year of the COVID-19 pandemic. In other hCoV species, we note that recombination has a predilection for structural genes and is relatively frequent in most medically significant hCoV over a relatively short evolution timescale. These findings are timely with the announcement of a more recent unpublished recombinant SARS-CoV-2 strain (25), as well as the recent focus on how insights can be learned from the functional implications of antigenic evolution now demonstrated in seasonal coronaviruses (26).

From more than 100,000 SARS-CoV-2 genomes, we noted 8 instances of recombination with moderate confidence. However, all were flagged by the RDP4 software as being possibly driven by other processes despite support by three or more recombination detection methods. This may reflect the relatively lower viral diversity across the first year of the pandemic. Similarly, we did not detect a high-confidence recombination signal in SARS-CoV-1, a virus with a limited temporal distribution. In contrast, we show that recombination was relatively frequent in seasonal coronavirus and MERS-CoV datasets comprising a longer period of sampling, including recombinants sufficiently fit for onward transmission. Furthermore, our analyses show that three of the coronaviruses (MERS, OC43 and NL63) had breakpoint propensity for structural genes. Several reasons for this may exist, such as inherent structural similarities of the coronaviruses in this region causing enhanced enzyme slippage, or positive selection pressure. The latter may be correlated with evasion of the human immune system.

Our endemic coronavirus analyses also highlighted that recombination affected the estimated temporal structure of coronavirus sequence datasets. This serves as a caution for genomic epidemiology studies which do not identify and account for recombinants in SARS-CoV-2 and other coronavirus analyses. Indeed, recombination has long been known to be a cause of phylogenetic incongruence for other viruses (23). Our time-scaled evolutionary analyses, adjusted for recombination to restore a molecular clock signal, yielded insights into the recent epidemiology of seasonal coronaviruses. We noted that the current sampled diversity of seasonal coronaviruses has emerged within a 70 year period, punctuated by new lineage emergence at intervals ranging from 5 to 20 years. Caution is required in inferring that these viruses spilled over into humans at these timepoints, however, as this may reflect the divergence of new lineages from prior, unsampled older viral populations circulating in humans. In the case of 229E hCoV, we noted that the full genome TMRCA estimates were preceded by the clinical reports of 229E infection in the 1960s (24) and a spike gene analysis incorporating older partial genome sequences from the earlier 20^th^ century showed lineage extinction and replacement. This is consistent with previous analyses suggesting that 229E evolution is characterized by prior lineage extinction and new lineage emergence (27).

It is important to note that our RNA genomic recombination detection in sequence data remains a statistical estimation only, as previously discussed in detail for dengue viruses (28). In addition, the size of our SARS-CoV-2 dataset was computationally prohibitive to perform recombination detection across all data, which might have resulted in missing additional recombination events. Also, unsampled data are a pervasive technical risk for recombination detection, as demonstrated by the variable finding of recombinant events across subsampled SARS-CoV2 data. Finally, our strict criteria for identifying recombination events with high confidence may have resulted in the removal of some true recombinant events.

Still, these findings provide critical insights into the possible projected evolution of SARS-CoV2, as it becomes an endemic viral pathogen in human populations. Recombination in other RNA viruses has been associated with changes in host tropism, virulence or epidemiology (1). Ongoing genomic-based COVID-19 surveillance has recently been highlighted as a critical public health tool to detect novel SARS-CoV-2 variants, such as the B.1.1.7, B.1.351, and P.1 variants (29, 30). Robust genomic surveillance will be essential for the timely detection of recombination in SARS-CoV-2, which may have implications for the diagnosis of and immunity to this pandemic pathogen.

## MATERIALS AND METHODS

### Data curation and alignment

Full genomes of endemic seasonal hCoVs, MERS-CoV and SARS-CoV-1 were downloaded from the NIAID Virus Pathogen Database and Analysis Resource (ViPR) (31, 32). Specifically, we obtained n = 22 hCoV-229E whole genomes, n = 138 hCoV-OC43 whole genomes, n = 68 hCoV-NL63 whole genomes, n = 37 HKU1 whole genomes, n = 365 MERS-CoV whole genomes (including human and camel host), and n = 49 SARS-CoV-1 whole genomes. We excluded laboratory constructs and sequences without host, collection date and location history. Datasets were aligned using MAFFT (33), with manual alignment thereafter in MEGA v6.0 (34). Alignment lengths were adjusted such that all genomes from one alignment were of approximately equal length (OC43 = 30,639 nt; NL63 = 27,483 nt; HKU1 = 29,892 nt; 229e = 27,292 nt; MERS = 29,985 nt; SARS1 = 29,719 nt). Partial genomes were excluded. Accession numbers for these data are presented in Supplemental Table 3.

All available full SARS-CoV-2 genomes (n = 157,439) up to October 23, 2020 were downloaded from the GISAID database (35). These data were curated by removing (i) any partial genomes (<90% full genome length), (ii) any genomes with > 100 continuous ambiguous base calls (Ns), and (iii) removing bat and pangolin non-human genomes. A single genome (Italy/CAM-IZSM-45946/2020) was removed due to the presence of non-IUPAC nucleotide codes. The remaining 100,296 genomes were aligned to the MN908947.3 reference using mafftc (7.471) in the MAFFT software (33). Alignment ends (first 265 and last 229 nt) were trimmed using the trimal command (36). This yielded a final alignment of length 29,409 bp.

### Recombination detection and determination of breakpoints

Datasets underwent recombination detection using the RDP4 software (37). Recombination signal detection was performed with a suite of methods: the original RDP method (38), BOOTSCAN (39), MAXCHI (40), CHIMAERA (41), 3SEQ (42), GENECONV (43), SISCAN (44). Following the detection of a ‘recombination signal’ with these methods, the approximate breakpoint positions were determined using a hidden Markov model, BURT, and the recombinant sequence identified using the PHYLPRO (45), VISRD (46), and EEEP methods (47).

For SARS-CoV2, due to the prohibitive computational demand of recombination detection analysis in a dataset of this size, we randomly subsampled, with replacement, n = 300 sequences with 100 x iterations (n = 30,000 full genomes) to perform recombination detection.

As individual recombination detection methods may have limited specificity, we developed a customized framework to ascertain the level of evidence for those recombination events detected in these datasets. Recombination events identified by only one or two methods in RDP4 were assigned a ‘low’ level of confidence, and those identified by at least three methods in RDP4 were assigned at least a ‘moderate’ level of confidence. We further assigned a ‘high’ level of confidence for those recombination events identified by at least three RDP4 methods with no other identified process which may have explained the recombination signal (37). The evolutionary processes that might be misinterpreted as recombination are typically caused by a combination of mutation rate variation between sites superimposed on mutation rate variation along lineages (M. Darren, personal communication, Dec 4, 2020). For those recombination events with a high level of evidence, the location of breakpoints were plotted across the whole genome and a χ^2^ test used to compare frequency of breakpoints at structural versus non-structural regions. Breakpoints that could not be placed with certainty due to overlapping recombination or other reasons were not included in the analyses.

### Estimation of temporal structure of hCoV with and without recombinant strains

Datasets for each hCoV species underwent nucleotide model substitution selection using JModelTest2 (48), with model selection as follows: 229E = SYM+I+G, HKU1 = GTR+G+I, NL63 = GTR+I, OC43 = GTR+G+I. A maximum likelihood phylogeny was inferred using the PhyML software (49), with aLRT for node support and tips labeled by date of collection. Root-to-tip regression was performed using the Temp-Est tool (50), with slope coefficient and intercept values used as preliminary estimates of evolutionary rate and time-to-most-common recent ancestor (TMRCA), respectively. Confirmed recombinant sequences were annotated on these phylogenies to identify recombinant clades and singletons. Phylogenetic analyses were repeated with the recombinants removed to estimate the impact of recombination on time-scales and temporal structure of hCoV evolution. For NL63, removal of recombinant genomes resulted in a small dataset and a small remaining fraction of the initial phylogenetic tree. Therefore, NL63 genomes were screened for a common region without any recombination signal. A region of 7105 nts (region 13093-20198 of the alignment) was found in which most genomes (n=56) had no recombination signal, and this region was used for subsequent time-scaled phylogenetic analyses, with the 9 genomes with recombination breakpoints in this region removed from the dataset.

### Estimating time-scales of emergence of the currently circulating seasonal hCoV lineages

We leveraged recombination-free endemic seasonal hCoV datasets, in addition to removing other root-to-tip regression outlier genomes (n = 3+4 identical genomes for OC43, n = 1 for NL63, n = 0 for 229e, n =2 for HKU1), to reconstruct the time-scale of emergence of currently circulating seasonal hCoV lineages across the 229E, OC43, NL63 and HKU1 types. We focused on these coronaviruses because they are well established viruses within human populations and may serve as a model for the projected evolutionary future of SARS-CoV-2 which is now already an endemic RNA virus. This is in contrast to the now extinct SARS-CoV-1 virus, and MERS-CoV, the latter which continues to be defined by more sporadic and discrete spillover events (51).

Time-scaled genealogies of these viruses were inferred using the BEAST software 1.8.4 (52). To minimize the risk of model misspecification, we inferred maximum clade credibility phylogenies with combinations of demographic models (constant, exponential and skyline population models) and molecular clock models (strict versus relaxed) (Supplemental Table 4). For each hCoV dataset, the optimal combination of demographic and molecular clock model was selected by logarithmic marginal likelihoods inferred by the path-sampling/ stepping-stone method (47).

## DISCLAIMER

Material has been reviewed by the Walter Reed Army Institute of Research. There is no objection to its presentation and/or publication. The View(s) expressed are those of the authors and do not necessarily reflect the official views of the Uniformed Services University of the Health Sciences, Henry M. Jackson Foundation for the Advancement of Military Medicine, Inc., Department of Health and Human Services, the National Institutes of Health, the Departments of the Army, Navy or Air Force, the Department of Defense, or the U.S. Government.

## ACKNOWLEDGEMENTS

The authors would like to acknowledge all the authors and originating and submitting laboratories of the sequences from GISAID’s EpiCov that were used in our analyses. A full acknowledgments list is submitted as Supplemental Table 5. This study was funded by the Global Emerging Infections Surveillance (GEIS) Branch (ProMIS ID: P0140_20_WR) and the US Department of Defense Health Agency. Support for this work was provided by the Infectious Disease Clinical Research Program (IDCRP), a Department of Defense program executed through the Uniformed Services University of the Health Sciences, Department of Preventive Medicine and Biostatistics through a cooperative agreement with The Henry M. Jackson Foundation for the Advancement of Military Medicine, Inc. (HJF).

