## Supplemental Table 1 for "A comparative recombination analysis of human coronaviruses and implications for the SARS-CoV-2 pandemic"

**Table S1. Recombinant events detected in SARS-CoV-2 dataset by RDP4 method positivity and genome frequency**

| Potential Recombinant Sequence* | Date | Data Subset[a] | Total number of sequences found with event | Positivity by RDP4 method | | | | | | | n methods positive total |
|---|---|---|---|---|---|---|---|---|---|---|---|
| | | | | RDP | GENE CONV | Bootscan | MaxChi | Chimaera | SiScan | 3Seq | |
| India/MH-NIV-218275/2020 | 9/19/2020 | 15 | 1 | no | no | no | yes | yes | no | no | 2 |
| Italy/LOM-UniMI02/2020 | 2/24/2020 | 21 | 1 | yes | yes | yes | yes | yes | no | yes | 6[b] |
| Australia/VIC10098/2020 | 8/12/2020 | 21 | 26 | yes | yes | yes | yes | yes | no | yes | 6[b] |
| India/MH-NIV-8199/2020 | 4/2/2020 | 23 | 1 | no | no | no | yes | yes | yes | no | 1 |
| India/GJ-GBRC71/2020 | 4/27/2020 | 29 | 166 | yes | no | yes | yes | no | yes | yes | 5[b] |
| USA/WA-UW-22852/2020 | 8/5/2020 | 29 | 1 | no | yes | no | no | no | no | no | 1 |
| England/QEUH-9B8E31/2020 | 9/7/2020 | 29 | 1 | no | yes | no | no | no | no | no | 1 |
| USA/WA-UW-10212/2020 | 5/31/2020 | 32 | 1 | no | no | no | no | no | no | yes | 1 |
| Switzerland/ZH-ETHZ-230030/2020 | 8/4/2020 | 40 | 1 | no | yes | no | no | no | no | no | 1 |
| Malaysia/190300/2020 | 3/22/2020 | 41 | 1 | yes | no | yes | yes | yes | no | yes | 5[b] |
| Germany/BE-RKI-Z-0006/2020 | 4/20/2020 | 41 | 2 | no | yes | no | no | no | no | no | 1 |
| USA/TX-HMH0430/2020 | 4/2/2020 | 42 | 154 | no | yes | no | no | no | no | no | 1 |
| USA/TX-HMH-6040/2020 | 7/1/2020 | 46 | 4 | no | yes | no | no | no | no | no | 1 |
| USA/MI-UM-10035994708/2020 | 9/17/2020 | 58 | 44 | no | no | no | yes | no | no | no | 1 |
| USA/WA-UW-12542/2020 | 6/27/2020 | 64 | 1 | no | no | no | no | no | no | yes | 1 |
| USA/TX-HMH-7945/2020 | 6/30/2020 | 65 | 1 | no | yes | no | no | no | no | no | 1 |
| England/QEUH-9B5371/2020 | 9/3/2020 | 65 | 1 | no | no | no | no | no | yes | yes | 2 |
| Scotland/CVR3535/2020 | 5/14/2020 | 70 | 1 | no | yes | no | no | no | no | no | 1 |
| USA/WI-GMF-14662/2020 | 8/10/2020 | 70 | 1 | no | yes | no | no | no | no | no | 1 |
| Turkey/6224-Ankara1034/2020 | 3/17/2020 | 73 | 1 | no | no | no | yes | no | no | no | 1 |
| England/20139052002/2020 | 3/27/2020 | 75 | 297 | yes | yes | yes | no | no | no | yes | 4[b] |
| USA/PR-B5XN/2020 | 8/11/2020 | 76 | 1 | no | yes | no | no | no | no | no | 1 |
| Turkey/ACUTG-1/2020 | 4/13/2020 | 78 | 1 | no | no | no | no | no | no | yes | 1 |

| | | | | | | | | | | | |
|---|---|---|---|---|---|---|---|---|---|---|---|
| England/201061439/2020 | 3/4/2020 | 82 | 296 | no | yes | no | no | no | no | no | 1 |
| USA/WA-UW-22859/2020 | 8/5/2020 | 85 | 1 | no | no | no | no | no | no | yes | 1 |
| USA/WA-UW-12771/2020 | 6/29/2020 | 86 | 151 | no | no | yes | yes | no | no | no | 2 |
| Wuhan/HBCDC-HB-04/2019 | 12/30/2019 | 87 | 1 | no | no | no | yes | no | no | yes | 2 |
| USA/WA-UW-19260/2020 | 8/1/2020 | 91 | 1 | no | no | no | no | no | no | yes | 1 |
| USA/TX-HMH-2673/2020 | 6/1/2020 | 93 | 5 | no | yes | no | no | no | no | no | 1 |
| India/KA-InStem-NCBS-0080/2020 | 6/17/2020 | 94 | 1 | yes | no | yes | no | yes | no | yes | 4[b] |
| England/MILK-99250B/2020 | 8/27/2020 | 94 | 4 | yes | no | yes | yes | no | no | yes | 4[b] |
| England/MILK-99A9BC/2020 | 8/29/2020 | 96 | 1 | no | no | no | no | no | no | yes | 1 |
| England/ALDP-9BDA83/2020 | 9/10/2020 | 99 | 264 | no | no | yes | yes | no | yes | yes | 4[b] |

[a]Refers to 100 subsets of data randomly subsampled (without replacement) from n = 100,296 SARS-CoV-2 full genomes

[b]Recombination event indicated to be possibly due to another process other than recombination (this is annotated for those events supported by ≥ 3 methods)

*SARS-CoV-2 sequence accession numbers available at https://github.com/conte1/recombination_in_coronaviruses
