## Supplemental Table 2 for "A comparative recombination analysis of human coronaviruses and implications for the SARS-CoV-2 pandemic"

**Table S2. Location of breakpoints in SARS-CoV-2 recombinants identified with moderate-level evidence***

| Potential Recombinant Sequence | Date | Data Subset[a] | Total number of sequences found with event | n methods positive total | Breakpoint locations | | |
|---|---|---|---|---|---|---|---|
| | | | | | Gene(s) involved | Begin (genome) | End (genome) |
| Australia/VIC10098/2020 | 8/12/2020 | 21 | 26 | 6 | nsp3 | 6983 | 7492 |
| Italy/LOM-UniMI02/2020 | 2/24/2020 | 21 | 1 | 6 | nsp3,nsp4 | 6498 | 8353 |
| India/GJ-GBRC71/2020 | 4/27/2020 | 29 | 166 | 5 | nsp3, nsp16 | 2586 | 25286 |
| Malaysia/190300/2020 | 3/22/2020 | 41 | 1 | 5 | nsp13, orf7a | 19728 | 27173 |
| England/20139052002/2020 | 3/27/2020 | 75 | 297 | 4 | nsp3, nsp4 | 6205 | 8688 |
| England/MILK-99250B/2020 | 8/27/2020 | 94 | 4 | 4 | spike_S1_RBD, spike_S2 | 22858 | 24152 |
| India/KA-InStem-NCBS-0080/2020 | 6/17/2020 | 94 | 1 | 4 | spike_S1_RBD | 22172 | 23072 |
| England/ALDP-9BDA83/2020 | 9/10/2020 | 99 | 264 | 4 | nsp2, ORF3a | 510 | 25409 |

*No recombinants were identified with high level evidence as all may have been explained by another process other than recombination
