## Supplemental Table 3 for "A comparative recombination analysis of human coronaviruses and implications for the SARS-CoV-2 pandemic"

**Table S3. hCoV Sequence Accession Number and Location/Date of Sampling**

| |
|---|
| **229E sequence data** |
| NA\|JX503060\|0349\|Netherlands\|2010 |
| NA\|JX503061\|J0304\|Italy\|2009 |
| NA\|KF514430\|229E/human/USA/933_50/1993\|USA\|1993_03_18 |
| NA\|KF514432\|229E/human/USA/932_72/1993\|USA\|1993_02_22 |
| NA\|KF514433\|229E/human/USA/933_40/1993\|USA\|1993_03_11 |
| NA\|KU291448\|HCoV_229E/BN1/GER/2015\|Germany\|2015 |
| NA\|KY369908\|HCoV_229E/Seattle/USA/SC579/2016\|USA\|2016 |
| NA\|KY369909\|HCoV_229E/Seattle/USA/SC677/2016\|USA\|2016 |
| NA\|KY369910\|HCoV_229E/Seattle/USA/SC1143/2016\|USA\|2016 |
| NA\|KY369911\|HCoV_229E/Seattle/USA/SC1212/2016\|USA\|2016 |
| NA\|KY369912\|HCoV_229E/Seattle/USA/SC9731/2016\|USA\|2016 |
| NA\|KY369913\|HCoV_229E/Seattle/USA/SC1073/2016\|USA\|2016 |
| NA\|KY369914\|HCoV_229E/Seattle/USA/SC9773/2016\|USA\|2016 |
| NA\|KY621348\|HCoV_229E/Seattle/USA/SC379/2016\|USA\|2016 |
| NA\|KY674914\|HCoV_229E/Seattle/USA/SC399/2016\|USA\|2016 |
| NA\|KY674919\|N08_434B\|USA\|2016 |
| NA\|KY684760\|HCoV_229E/Seattle/USA/SC2282/2017\|USA\|2016 |
| NA\|KY967357\|HCoV_229E/Seattle/USA/SC2872/2015\|USA\|2015 |
| NA\|KY983587\|HCoV_229E/Seattle/USA/SC3112/2015\|USA\|2015 |
| NA\|MF542265\|229E/Haiti_1/2016\|Haiti\|2016_03 |
| NA\|MN306046\|HCoV_229E/Seattle/USA/SC0865/2019\|USA\|2019 |
| NA\|MN369046\|HCoV_229E/Seattle/USA/SC9724/2018\|USA\|2018 |
| **HKU1 sequence data** |
| BJ01_p3\|Human\|KT779555\|NA\|China\|2009-04-23 |
| BJ01_p9\|Human\|KT779556\|NA\|China\|2009-04-23 |
| Caen1\|Human\|HM034837\|NA\|France\|2005-03-04 |
| HKU1/human/USA/1102/2005\|Human\|KF850450\|NA\|USA\|2005-04-09 |

HKU1/human/USA/HKU1_4/2005|Human|KF430196|NA|USA|2005-04-19
HKU1/human/USA/HKU1_14/2009|Human|KF430199|NA|USA|2009-12-28
HKU1/human/USA/HKU1_20/2010|Human|KF686345|NA|USA|2010-01-30
HKU1|Human|AY597011|NA|China|2004-1
HKU1|Human|NC_006577|NA|China|2004-1
KF430200|NA|USA|2010-01-08
KF430201|NA|USA|2010-01-22
KF430202|NA|USA|2010-01-03
KF686338|NA|USA|2005-02-18
KF686339|NA|USA|2009-10-31
KF686340|NA|USA|2009-11-28
KF686341|NA|USA|2010-01-16
KF686342|NA|USA|2009-12-13
KF686343|NA|USA|2010-01-08
KF686344|NA|USA|2009-12-28
KF686346|NA|USA|2010-01-09
MH940245|NA|Thailand|2017-06-04
N3|Unknown|DQ415903|NA|China|2003-04
N6|Unknown|DQ415904|NA|China|2004-01
N7|Unknown|DQ415905|NA|China|2004-01
N08_87|Human|KY674921|NA|USA|2016
N09_1605B|Human|KY674943|NA|USA|2016
N09_1627B|Human|KY674942|NA|USA|2016
N09_1663B|Human|KY674941|NA|USA|2016
N9|Unknown|DQ415906|NA|China|2004-03
N10|Unknown|DQ415907|NA|China|2004-03
N11|Unknown|DQ415908|NA|Hong_Kong|2004-04
N13|Unknown|DQ415909|NA|China|2004-05
N14|Unknown|DQ415910|NA|Hong_Kong|2004-07
N15|Unknown|DQ415911|NA|China|2004-11
N16|Unknown|DQ415912|NA|China|2004-11
N17|Unknown|DQ415913|NA|China|2004-11

N18|Unknown|DQ415914|NA|China|2004-11
N19|Unknown|DQ415896|NA|Hong_Kong|2004-12
N20|Unknown|DQ415897|NA|Hong_Kong|2004-12
N21|Unknown|DQ415898|NA|Hong_Kong|2004-12
N22|Unknown|DQ415899|NA|China|2005-01
N23|Unknown|DQ415900|NA|China|2005-02
N24|Unknown|DQ415901|NA|China|2005-01
N25|Unknown|DQ415902|NA|China|2005-02
SC2521|Human|MK167038|NA|USA|2017

**NL63 sequence data**

Human_coronavirus_NL63|UNKNOWN_AY518894|AY518894|Netherlands|1988_04
Human_coronavirus_NL63|Amsterdam_496|DQ445912|Netherlands|2003_02
Human_coronavirus_NL63|NL63/DEN/2005/232|JQ765569|USA|2005_01_18
Human_coronavirus_NL63|NL63/DEN/2005/235|JQ765570|USA|2005_01_19
Human_coronavirus_NL63|NL63/human/USA/0111_25/2001|KF530112|USA|2001_11_21
Human_coronavirus_NL63|NL63/DEN/2009/9|JQ765563|USA|2009_03_16
Human_coronavirus_NL63|NL63/DEN/2009/14|JQ765564|USA|2009_03_01
Human_coronavirus_NL63|NL63/DEN/2009/15|JQ765565|USA|2009_02_13
Human_coronavirus_NL63|NL63/DEN/2009/22|JQ900256|USA|2009_03_03
Human_coronavirus_NL63|NL63/DEN/2005/291|JQ900258|USA|2005_01_26
Human_coronavirus_NL63|CN0601/14|MG772808|South_Korea|2014_11_25
Human_coronavirus_NL63|ChinaGD04|MK334047|China|2018_07_25
Human_coronavirus_NL63|ChinaGD01|MK334046|China|2018_09_01
Human_coronavirus_NL63|Amsterdam_057|DQ445911|Netherlands|2002_12
Human_coronavirus_NL63|HCoV_NL63/Seattle/USA/SC2940/2015|KY983586|USA|2015
Human_coronavirus_NL63|CBJ_037|JX104161|China|2008_11_04
Human_coronavirus_NL63|CBJ123|JX524171|China|2009_01_24
Human_coronavirus_NL63|NL63/DEN/2009/31|JQ900257|USA|2009_02_21
Human_coronavirus_NL63|Kilifi_HH_5709_19_May_2010|MG428699|Kenya|2010_05_19
Human_coronavirus_NL63|Kilifi_HH_0512_04_Jun_2010|MG428701|Kenya|2010_06_04
Human_coronavirus_NL63|Kilifi_HH_0511_01_Jun_2010|MG428703|Kenya|2010_06_01
Human_coronavirus_NL63|Kilifi_HH_5402_20_May_2010|MG428704|Kenya|2010_05_20
Human_coronavirus_NL63|HCoV_NL63/Haiti_1/2015|KT266906|Haiti|2015_01_16
Human_coronavirus_NL63|Kilifi_HH_3807_11_May_2010|MG428702|Kenya|2010_05_11
Human_coronavirus_NL63|Kilifi_HH_3808_24_May_2010|MG428706|Kenya|2010_05_24
Human_coronavirus_NL63|Kilifi_HH_0522_21_May_2010|MG428705|Kenya|2010_05_21

Human_coronavirus_NL63|HCoV_NL63/Seattle/USA/SC0768/2019|MN306040|USA|2019
Human_coronavirus_NL63|Amsterdam_I|AY567487|Netherlands|2003_01
Human_coronavirus_NL63|Amsterdam_I|NC_005831|Nehterlands|2003_01
Human_coronavirus_NL63|NL63/RPTEC/2004|JX504050|USA|2004
Human_coronavirus_NL63|UNKNOWN_DJ009246|DJ009246|Netherlands|2003_01
Human_coronavirus_NL63|UNKNOWN_CS124012|CS124012|Netherlands|2003_01
Human_coronavirus_NL63|NL63/DEN/2005/1062|JQ765573|USA|2005_04_12
Human_coronavirus_NL63|NL63/DEN/2005/193|JQ765568|USA|2005_01_11
Human_coronavirus_NL63|NL63/DEN/2005/449|JQ900259|USA|2005_02_09
Human_coronavirus_NL63|NL63/DEN/2005/1876|JQ765575|USA|2005_11_21
Human_coronavirus_NL63|NL63/DEN/2005/271|JQ765571|USA|2005_01_23
Human_coronavirus_NL63|NL63/DEN/2005/347|JQ765572|USA|2005_02_01
Human_coronavirus_NL63|NL63/DEN/2005/1862|JQ765574|USA|2005_11
Human_coronavirus_NL63|N07_196B|KY554969|USA|2016
Human_coronavirus_NL63|N07_262B|KY829118|USA|2015
Human_coronavirus_NL63|N07_468B_176X|KY554971|USA|2016
Human_coronavirus_NL63|NL63/human/USA/012_31/2001|KF530105|USA|2001_02_23
Human_coronavirus_NL63|NL63/DEN/2008/16|JQ765566|USA|2008_01_08
Human_coronavirus_NL63|NL63/DEN/2009/6|JQ900255|USA|2009_02_25
Human_coronavirus_NL63|N07_324B_182X|KY554970|USA|2016
Human_coronavirus_NL63|N07_6B|KY674915|USA|2016
Human_coronavirus_NL63|N06_1144B|KY554967|USA|2016
Human_coronavirus_NL63|N07_185B|KY554968|USA|2016
Human_coronavirus_NL63|N07_64B|KY674916|USA|2016
Human_coronavirus_NL63|NL63/DEN/2009/20|JQ765567|USA|2009_03_12
Human_coronavirus_NL63|NL63/UF_2/2015|KU521535|USA|2015_09_01
Human_coronavirus_NL63|NL63/UF_1/2015|KT381875|USA|2015
Human_coronavirus_NL63|NL63/UF_2/2015|KX179500|USA|2015_09
Human_coronavirus_NL63|NL63/DEN/2005/1120|JQ900260|USA|2005_04_25
Human_coronavirus_NL63|NL63/human/USA/891_6/1989|KF530108|USA|1989_01_05
Human_coronavirus_NL63|NL63/human/USA/891_4/1989|KF530114|USA|1989_01_03
Human_coronavirus_NL63|NL63/human/USA/904_20/1990|KF530104|USA|1990_04_26
Human_coronavirus_NL63|NL63/human/USA/903_28/1990|KF530109|USA|1990_03_21
Human_coronavirus_NL63|NL63/human/USA/901_24/1990|KF530111|USA|1990_01_05
Human_coronavirus_NL63|NL63/human/USA/905_25/1990|KF530113|USA|1990_05_29
Human_coronavirus_NL63|NL63/human/USA/911_56/1991|KF530107|USA|1991_01_24
Human_coronavirus_NL63|NL63/human/USA/838_9/1983|KF530110|USA|1983_08_16
Human_coronavirus_NL63|HCoV_NL63/Seattle/USA/SC0179/2018|MN306018|USA|2018

| |
|---|
| Human_coronavirus_NL63\|ChinaGD02\|MK334043\|China\|2018_08_15 |
| Human_coronavirus_NL63\|ChinaGD03\|MK334044\|China\|2018_07_13 |
| Human_coronavirus_NL63\|ChinaGD05\|MK334045\|China\|2018_08_14 |
| Human_coronavirus_NL63\|NL63/human/USA/8712_17/1987\|KF530106\|USA\|1987_12_16 |
| NA\|MN369046\|HCoV_229E/Seattle/USA/SC9724/2018\|USA\|2018 |
| **OC43 sequence data** |
| NA\|KF923918\|10108/2010\|China\|2010_05 |
| NA\|KF923922\|8164/2009\|China\|2009_03 |
| NA\|KF923925\|10574/2010\|China\|2010_09 |
| NA\|KJ958218\|LY341\|China\|2011_10_03 |
| NA\|KJ958219\|LY342\|China\|2011_10_04 |
| NA\|KF923890\|39A/2007\|China\|2007_04 |
| NA\|KF923907\|5370/2007\|China\|2007_05 |
| NA\|KF923911\|5479/2007\|China\|2007_06 |
| NA\|KF923914\|5508/2007\|China\|2007_06 |
| NA\|KF923912\|5484/2007\|China\|2007_06 |
| NA\|KF923909\|5442/2007\|China\|2007_06 |
| NA\|KF923901\|5472/2007\|China\|2007_06 |
| NA\|KF923919\|5595/2007\|China\|2007_07 |
| NA\|KF923910\|5445/2007\|China\|2007_06 |
| NA\|KF923892\|5345/2007\|China\|2007_05 |
| NA\|KF923920\|5617/2007\|China\|2007_07 |
| NA\|KF923913\|5485/2007\|China\|2007_06 |
| NA\|KF923915\|5517/2007\|China\|2007_06 |
| NA\|KF923891\|5240/2007\|China\|2007_05 |
| NA\|KF923894\|5352/2007\|China\|2007_05 |
| NA\|KF923917\|5566/2007\|China\|2007_06 |
| NA\|KY554974\|N08_33B_360X\|USA\|2016 |
| NA\|KY554975\|N09_382B\|USA\|2016 |
| NA\|KY674920\|N09_595B\|USA\|2016 |
| NA\|KF923916\|5519/2007\|China\|2007_06 |
| NA\|KY554972\|N07_1541B_433X\|USA\|2016 |
| NA\|KY554973\|N07_1689B_116X\|USA\|2016 |
| NA\|KY674918\|N07_1647B\|USA\|2016 |
| NA\|KY674917\|N07_1609B\|USA\|2016 |
| NA\|KF923921\|69A/2007\|China\|2007_05 |
| NA\|KF923908\|5414/2007\|China\|2007_06 |

NA|KF923923|892A/2008|China|2008_10
NA|KF923893|2151A/2010|China|2010_07
NA|KF923924|10290/2010|China|2010_07
NA|KX344031|OC43/human/Mex/LRTI_238/2011|Mexico|2011_02_09
D|JN129835|HK04_02|China|2004_11
NA|KF923903|12691/2012|China|2012_05
NA|KX538977|MY_U1140/12|Malaysia|2012_09_10
NA|KX538969|MY_U523/12|Malaysia|2012_05_18
NA|KX538968|MY_U464/12|Malaysia|2012_05_09
NA|KX538971|MY_U732/12|Malaysia|2012_06_25
NA|KX538973|MY_U868/12|Malaysia|2012_07_16
NA|KX538974|MY_U945/12|Malaysia|2012_08_01
NA|KX538965|MY_U208/12|Malaysia|2012_03_28
NA|KX538967|MY_U413/12|Malaysia|2012_05_02
NA|KX538975|MY_U1024/12|Malaysia|2012_08_24
NA|KF923897|3269A/2012|China|2012_06
C|JN129834|HK04_01|China|2004_11
NA|KF923905|229/2005|China|2005_06
NA|KF923899|3582/2006|China|2006_09
NA|KF923900|3647/2006|China|2006_10
NA|KF923902|12689/2012|China|2012_05
NA|KF923904|12694/2012|China|2012_05
NA|KX538964|MY_U002/12|Malaysia|2012_02_22
NA|KX538976|MY_U1057/12|Malaysia|2012_08_27
NA|KX538972|MY_U774/12|Malaysia|2012_07_04
NA|KX538966|MY_U236/12|Malaysia|2012_04_02
NA|KX538970|MY_U710/12|Malaysia|2012_06_20
NA|KX538978|MY_U1758/13|Malaysia|2013_01_02
NA|KX538979|MY_U1975/13|Malaysia|2013_02_15
NA|KY967356|HCoV_OC43/Seattle/USA/SC2924/2015|USA|2015
NA|MG977451|TNP_12636|Cote_dIvoire|2016_12_10
NA|MG977452|TNP_12643|Cote_dIvoire|2016_12_10
NA|MF374983|HCoV_OC43/USA/TCNP_0070/2016|USA|2016_02_01
NA|MH121121|HCoV_OC43/USA/ACRI_0213/2016|USA|2016_12_19
NA|KY369907|HCoV_OC43/Seattle/USA/SC9741/2016|USA|2016
NA|KY983588|HCoV_OC43/Seattle/USA/SC3118/2015|USA|2015
NA|KY369905|HCoV_OC43/Seattle/USA/SC831/2016|USA|2016
NA|KY369906|HCoV_OC43/Seattle/USA/SC622/2016|USA|2016

NA|KY684759|HCoV_OC43/Seattle/USA/SC2269/2016|USA|2016
NA|KY967361|HCoV_OC43/Seattle/USA/SC2345/2015|USA|2015
NA|KY983583|HCoV_OC43/Seattle/USA/SC2481/2015|USA|2015
NA|KY967358|HCoV_OC43/Seattle/USA/SC2770/2015|USA|2015
NA|KY983585|HCoV_OC43/Seattle/USA/SC2854/2015|USA|2015
NA|MF374985|HCoV_OC43/USA/TCNP_00212/2017|USA|2017_01_17
NA|KY967359|HCoV_OC43/Seattle/USA/SC2730/2015|USA|2015
NA|MF374984|HCoV_OC43/USA/TCNP_00204/2017|USA|2017_01_03
NA|MN306036|HCoV_OC43/Seattle/USA/SC0682/2019|USA|2019
NA|MN306041|HCoV_OC43/Seattle/USA/SC0810/2019|USA|2019
NA|MN306042|HCoV_OC43/Seattle/USA/SC0839/2019|USA|2019
NA|MN306053|HCoV_OC43/Seattle/USA/SC9430/2018|USA|2019
NA|MN310478|HCoV_OC43/Seattle/USA/SC0776/2019|USA|2019
NA|MN026164|OC43_KLF_01_2018|Kenya|2018_01_18
NA|KF923895|10285/2010|China|2010_07
NA|KF530068|OC43/human/USA/007_11/2000|USA|2000_07_27
NA|KF530081|OC43/human/USA/991_5/1999|USA|1999_01_07
NA|KF530070|OC43/human/USA/991_19/1999|USA|1999_01_15
NA|KF530063|OC43/human/USA/9612_48/1996|USA|1996_12_30
NA|KF530099|OC43/human/USA/971_5/1997|USA|1997_01_02
NA|KF530069|OC43/human/USA/982_4/1998|USA|1998_02_05
NA|KF530088|OC43/human/USA/901_54/1990|USA|1990_01_23
NA|KF530071|OC43/human/USA/925_1/1992|USA|1992_05_04
NA|KF530076|OC43/human/USA/911_11/1991|USA|1991_01_03
NA|KF530091|OC43/human/USA/911_58/1991|USA|1991_01_24
NA|KF530089|OC43/human/USA/911_66/1991|USA|1991_01_29
NA|KF530082|OC43/human/USA/912_11/1991|USA|1991_02_07
NA|KF530094|OC43/human/USA/912_36/1991|USA|1991_02_22
NA|KF530079|OC43/human/USA/913_29/1991|USA|1991_03_14
NA|KF530067|OC43/human/USA/912_10/1991|USA|1991_02_07
NA|KF530096|OC43/human/USA/911_38/1991|USA|1991_01_15
NA|KF530095|OC43/human/USA/912_6/1991|USA|1991_02_05
NA|KF530084|OC43/human/USA/951_18/1995|USA|1995_01_12
NA|KF530098|OC43/human/USA/965_6/1996|USA|1996_05_10
NA|KF923886|1908A/2010|China|2010_03
NA|KF923889|1926/2006|China|2006_03
NA|KF923887|1997A/2010|China|2010_04
NA|KF923888|2145A/2010|China|2010_07

NA|KF923898|3184A/2012|China|2012_03
NA|MN306043|HCoV_OC43/Seattle/USA/SC0841/2019|USA|2019
NA|MN310476|HCoV_OC43/Seattle/USA/SC9428/2018|USA|2019
NA|KP198611|1783A/10|China|2010_01
NA|KP198610|2058A/10|China|2010_06
NA|KY014282|2007_09|France|2007
NA|MF314143|HCoV_OC43/USA/ACRI_0052/2016|USA|2016_03_07
NA|KY967360|HCoV_OC43/Seattle/USA/SC2476/2015|USA|2015
NA|KU131570|HCoV_OC43/UK/London/2011|United_Kingdom|2011_08_20
NA|KF923896|3074A/2012|China|2012_02
NA|KF923906|3194A/2012|China|2012_03
NA|KY014281|2002_04|France|2002
NA|KF530092|OC43/human/USA/008_5/2000|USA|2000_08_08
NA|KF530078|OC43/human/USA/9612_29/1996|USA|1996_12_17
NA|KF530064|OC43/human/USA/9612_9/1996|USA|1996_12_04
NA|KF530072|OC43/human/USA/9712_13/1997|USA|1997_12_11
NA|KF530080|OC43/human/USA/9712_31/1997|USA|1997_12_18
NA|KF530060|OC43/human/USA/851_15/1985|USA|1985_01_08
NA|KF530086|OC43/human/USA/872_5/1987|USA|1987_02_10
NA|KF530083|OC43/human/USA/873_19/1987|USA|1987_03_17
NA|KF530077|OC43/human/USA/873_16/1987|USA|1987_03_12
NA|KF530087|OC43/human/USA/873_6/1987|USA|1987_03_05
NA|KF530085|OC43/human/USA/871_25/1987|USA|1987_01_22
NA|KF530073|OC43/human/USA/8912_37/1989|USA|1989_12_21
NA|KF530065|OC43/human/USA/901_41/1990|USA|1990_01_17
NA|KF530066|OC43/human/USA/901_33/1990|USA|1990_01_16
NA|KF530061|OC43/human/USA/901_43/1990|USA|1990_01_19
NA|KF530097|OC43/human/USA/9211_43/1992|USA|1992_11_30
NA|KF530090|OC43/human/USA/931_85/1993|USA|1993_01_26
NA|KF530074|OC43/human/USA/9212_33/1992|USA|1992_12_16
NA|KF530075|OC43/human/USA/953_23/1995|USA|1995_03_09
