## Supplemental Table 4 for "A comparative recombination analysis of human coronaviruses and implications for the SARS-CoV-2 pandemic"

**Table S4. Model fit for comparative TMCRA estimates of 229E, NL63, OC43, and HKU1 hCoV**

| hCoV type | Clock model | Demographic model | Log marginal likelihood | MCMC chain |
|---|---|---|---|---|
| 229E WG | Strict | Constant | -42436.48 | 200 mil |
| 229E WG | Strict | Exponential | 1711388.9 | 200 mil |
| 229E WG | Strict | Skyline | 1141416.68 | 100 mil |
| 229E WG | UCLN | Constant | -42878.79 | 100 mil |
| 229E WG | UCLN | Exponential | -42879.04 | 100 mil |
| 229E WG | UCLN | Skyline | -42873.31 | 200 mil |
| 229E RBD S1 gene | UCLN | Skyline | _ | 600 mil |
| 229E S gene | UCLN | Skyline | _ | 600 mil |
| 229E N gene | UCLN | Skyline | _ | 600 mil |
| 229E N and S gene | UCLN | Skyline | _ | 600 mil |
| OC43 WG[a] | Strict | Constant | -59899.62 | 600 mil |
| OC43 WG[a] | Strict | Exponential | -59895.68 | 600 mil |
| OC43 WG[a] | Strict | Skyline | -59881.69 | 600 mil |
| OC43 WG[a] | UCLN | Constant | -59863.16 | 600 mil |
| OC43 WG[a] | UCLN | Exponential | -59866.145 | 600 mil |
| OC43 WG[a] | UCLN | Skyline | -59851.46 | 600 mil |
| NL63 WG[a] | Strict | Constant | -42203 | 600 mil |
| NL63 WG[a] | Strict | Exponential | -42206.51 | 600 mil |
| NL63 WG[a] | Strict | Skyline | -42203.81 | 600 mil |
| NL63 WG[a] | UCLN | Constant | -42194.37 | 600 mil |
| NL63 WG[a] | UCLN | Exponential | -42199.37 | 600 mil |
| NL63 WG[a] | UCLN | Skyline | -42188.97 | 600 mil |
| NL63[b] | Strict | Constant | -11745.53 | 600 mil |
| NL63[b] | Strict | Exponential | -11747.46 | 600 mil |
| NL63[b] | Strict | Skyline | -11746.42 | 600 mil |
| NL63[b] | UCLN | Constant | infinity | 600 mil |

| | | | | |
|---|---|---|---|---|
| NL63[b] | UCLN | Exponential | infinity | 600 mil |
| NL63[b] | UCLN | Skyline | -11743.43 | 600 mil |
| HKU1 WG[a] | Strict | Constant | -51641.8511 | 600 mil |
| HKU1 WG[a] | Strict | Exponential | -51641.6057 | 600 mil |
| HKU1 WG[a] | Strict | Skyline | -51523.685 | 600 mil |
| HKU1 WG[a] | UCLN | Constant | -51496.93 | 600 mil |
| HKU1 WG[a] | UCLN | Exponential | -51498.577 | 600 mil |
| HKU1 WG[a] | UCLN | Skyline | -51495.71 | 600 mil |

UCLN = uncorrelated lognormal (relaxed clock)

WG = whole genome

[a]Recombinant genomes removed

[b]Recombinant section removed
