## Supplementary figures and images for "A comparative recombination analysis of human coronaviruses and implications for the SARS-CoV-2 pandemic"

### Supplemental Figure 5

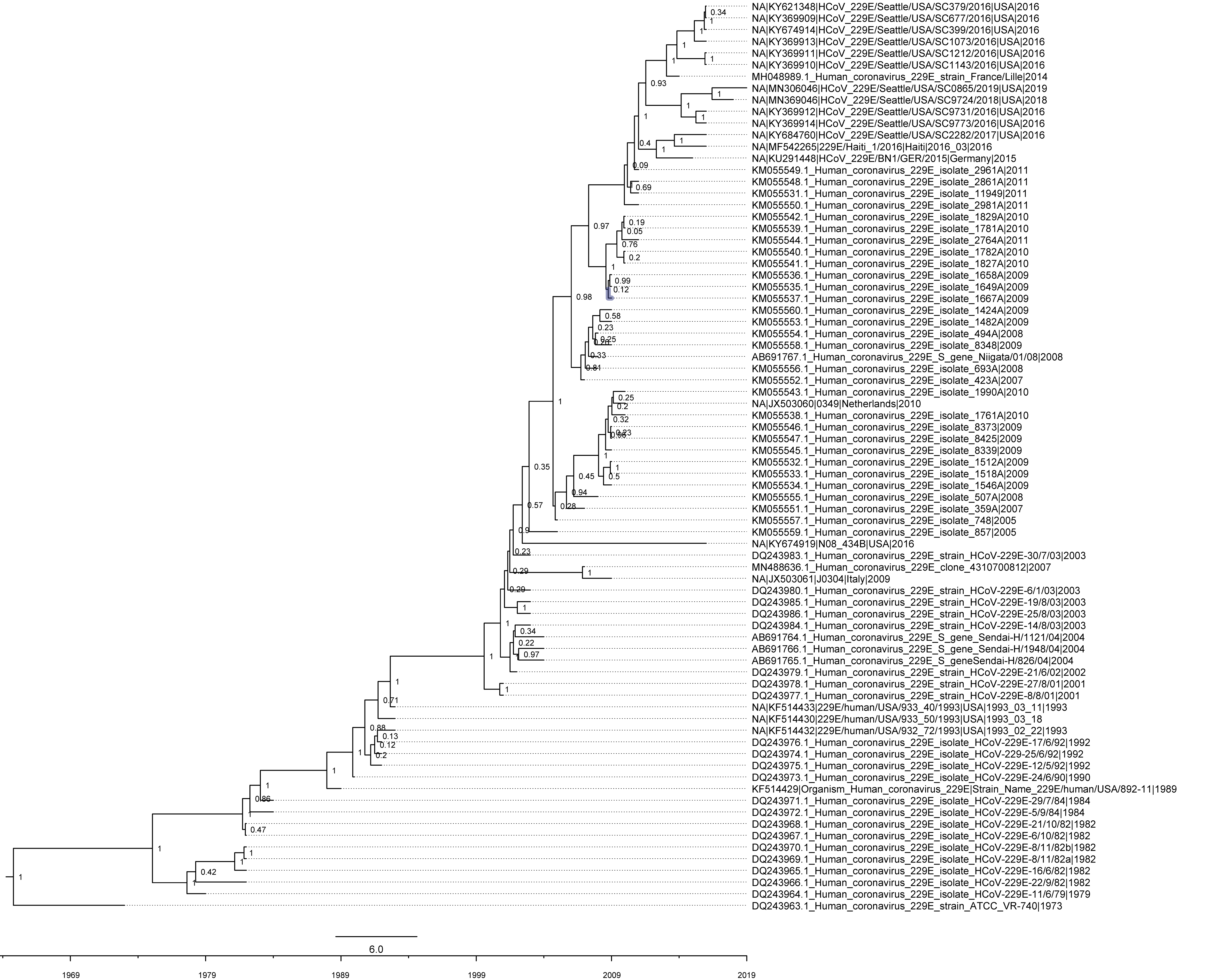

### Supplemental Figures 1-4

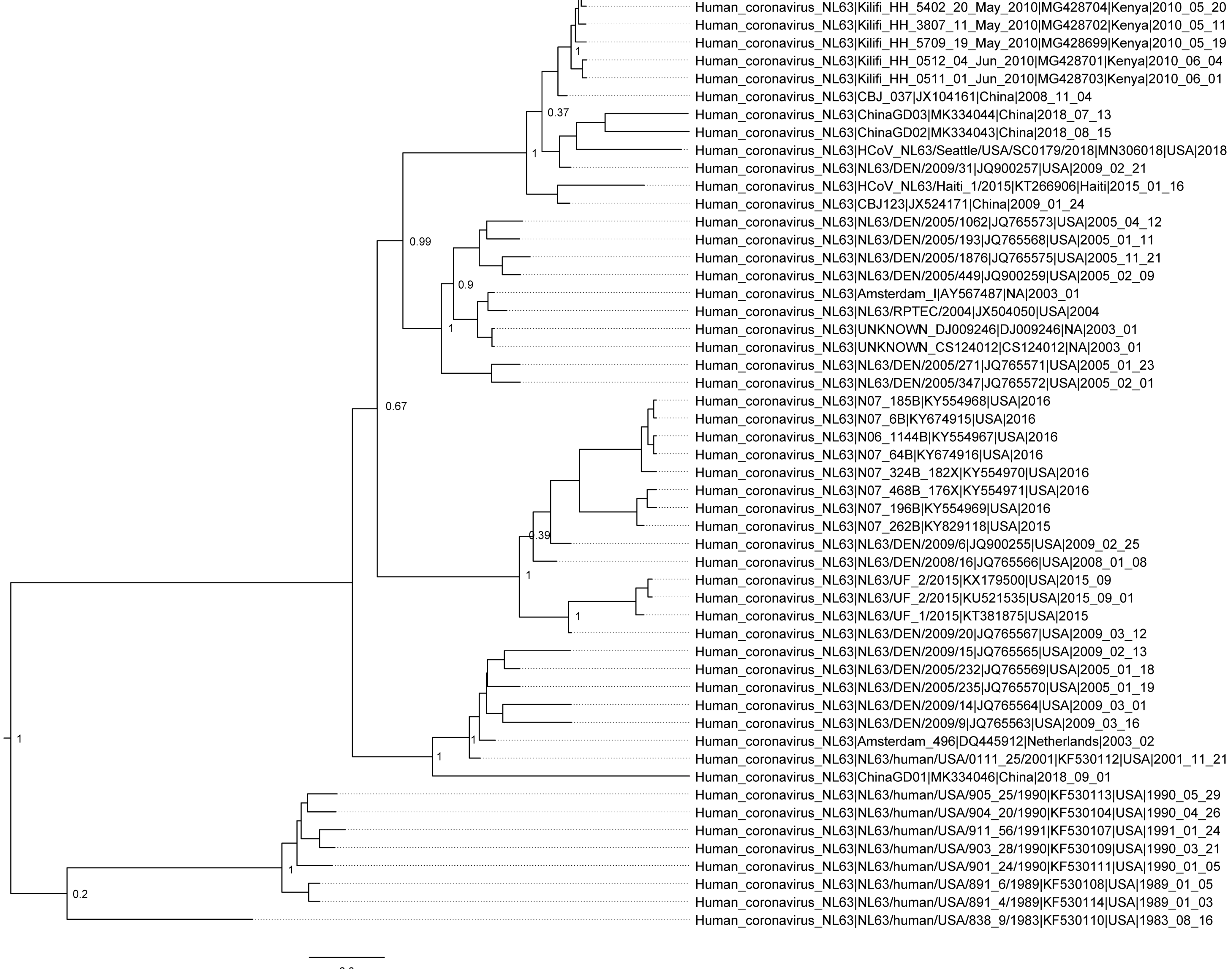

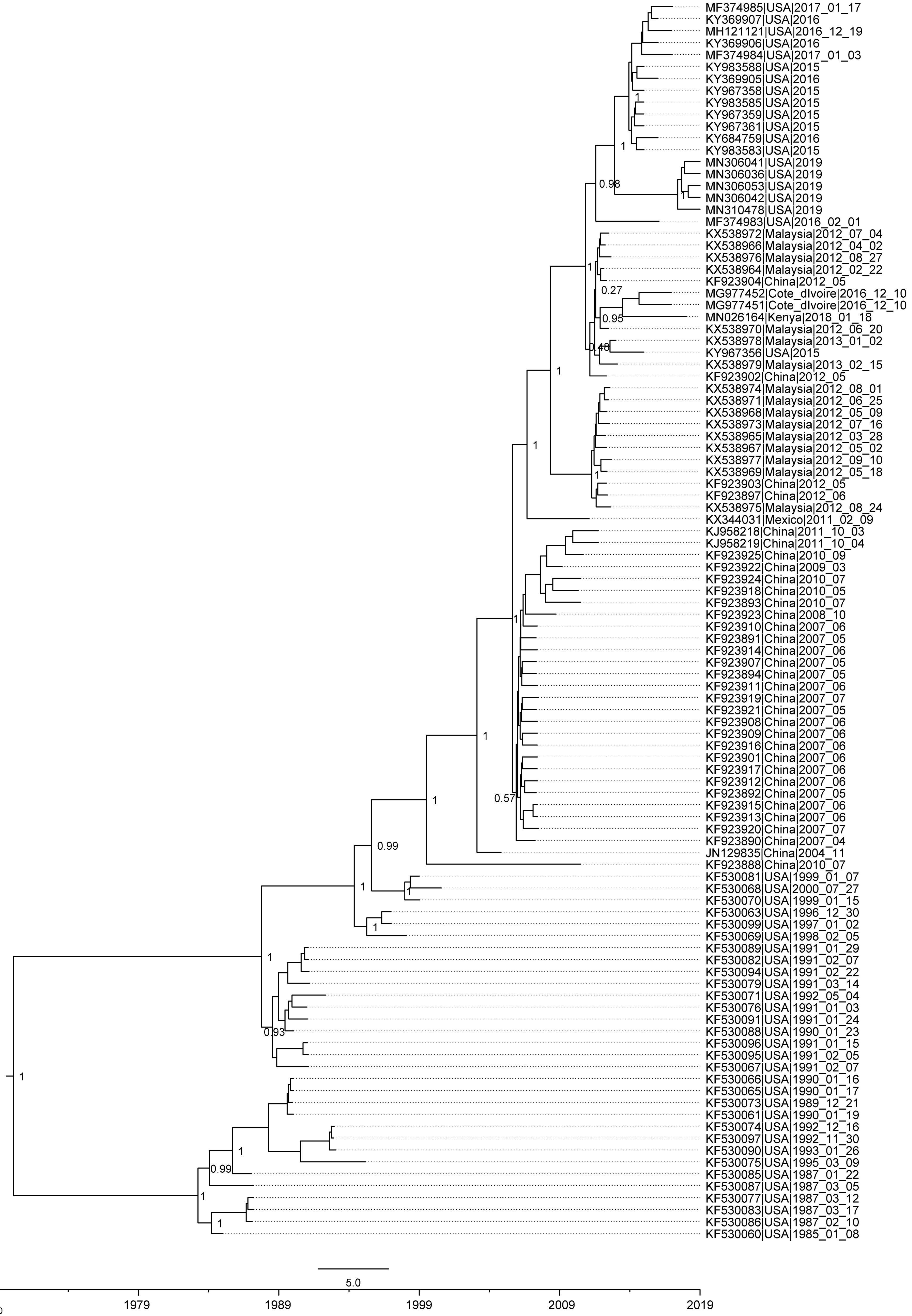

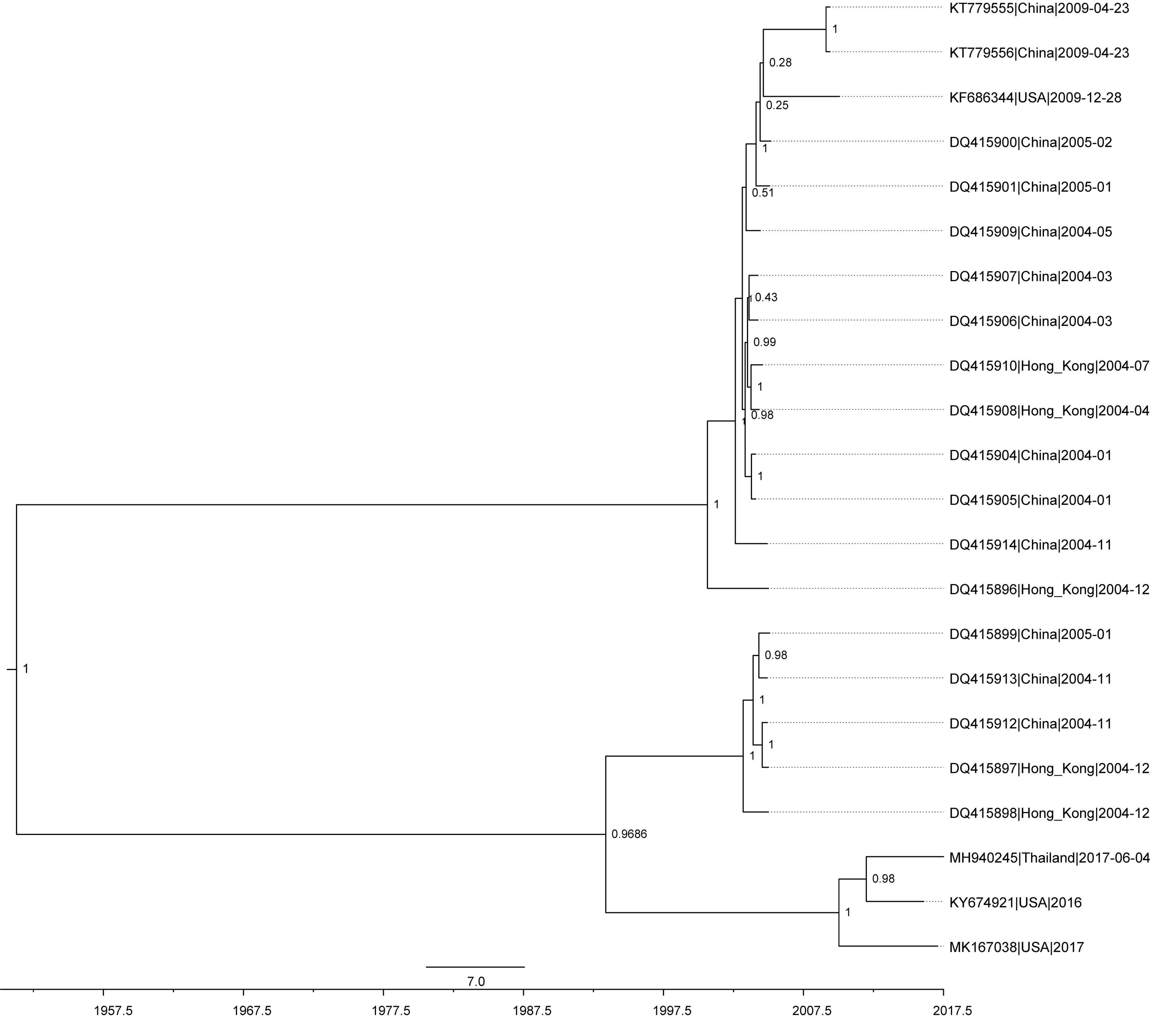

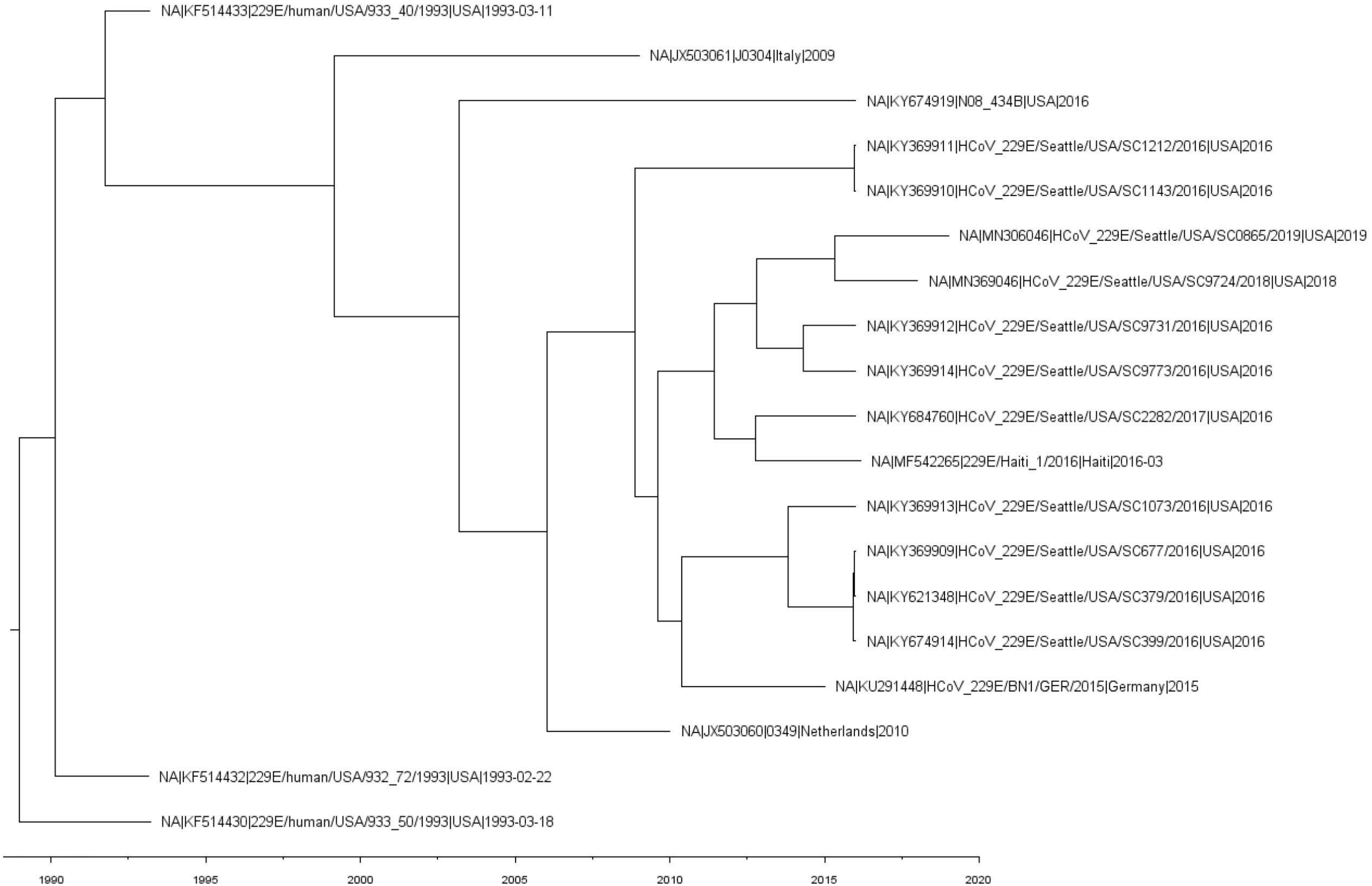
